# Structure of the energy converting methyltransferase (Mtr) of *Methanosarcina maze*i in complex with an oxygen-stress responsive small protein

**DOI:** 10.1101/2025.06.02.657420

**Authors:** Tristan Reif-Trauttmansdorff, Eva Herdering, Stefan Bohn, Tomas Pascoa, Jörg Kahnt, Erik Zimmer, Anuj Kumar, Ruth A. Schmitz, Jan M. Schuller

## Abstract

Methanogenic archaea contribute 1–2 Gt of methane annually, impacting both global carbon cycling and climate. Central to their energy metabolism is a membrane-bound, sodium-translocating methyltransferase complex: the N⁵-tetrahydromethanopterin:CoM-S-methyltransferase (Mtr complex), which catalyzes the methyl transfer between two methanogen specific cofactors. This exergonic methyl transfer step is coupled with a vectorial sodium ion transport from the cytoplasm to the cell exterior and is the only energy conserving step in hydrogenotrophic methanogenesis. Here, we present a 2.1 Å single-particle cryo-EM structure of the full Mtr complex from *Methanosarcina mazei*. Our structural model encompasses the entire complex, reveals the arrangement of archaeal phospholipids, the architecture of the sodium ion binding site, and the structure and interactions of all catalytic subunits. Most strikingly, we discover and characterize MtrI, a previously unannotated small open reading frame (small ORF), encoded protein (<100 aa) conserved across the order of Methanosarcinales. MtrI binds to the cytoplasmic domain of MtrA in response to oxygen exposure, suggesting a role in oxygen stress response and protection. By binding on top of the sodium channel and anchoring to the cobamide cofactor in MtrA’s cytoplasmic domain, MtrI might prevent sodium leakage and inhibit MtrA-CoM turnover. These findings offer new insights into methanogen energy conservation and uncover a potential adaptive response to oxygen exposure, expanding our understanding of methanogen survival strategies under oxidative stress.

## Introduction

Methanogenic archaea are widespread in different environments with high abundancies growing under strictly anaerobic conditions and are crucial for the last step in anaerobic degradation of organic matter. They represent the only organisms to produce methane as a catabolic end product of their energy metabolism. Consequently, they produce approximately one gigaton of this climate relevant gas (greenhouse gas) per year (reviewed in Thauer, 1998). There are several, phylogenetically not related pathways of methanogenesis using different substrates: e.g., hydrogenotrophic methanogenesis (H_2_/CO_2_), acetoclastic methanogenesis (acetate) and methylotrophic methanogenesis (e.g. methylamines and methanol) (Deppenmeier et al., 2002, Kurth, Op den Camp and Welte, 2020). A pivotal step in all methanogenic routes is catalyzed by the membrane-bound N⁵-methyltetrahydromethanopterin:coenzyme M methyltransferase complex (Mtr), which mediates the exergonic transfer of a methyl group from methyl-tetrahydromethanopterin (methyl-H₄MPT) or methyl-tetrahydrosarcinapterin (H₄SPT in some species) to HS-CoM (ΔG°′ = −30 kJ/mol), coupled to sodium ion translocation across the membrane (Müller, Blaut and Gottschalk, 1988; Gärtner *et al*., 1993; Weiss, Gärtner and Thauer, 1994; Lienard *et al*., 1996). This sodium-motive force drives chemiosmotic energy conservation and ATP synthesis via a Na⁺-dependent ATP synthase, forming the basis of what is often referred to as sodium-based bioenergetics in methanogenic archaea. In methylotrophic methanogens that disproportionate methanol into methane and CO₂, this reaction is reversed: the sodium ion gradient is used to drive the endergonic methyl transfer from methyl-CoM back to H₄MPT. Similarly, anaerobic methanotrophic archaea (ANME) oxidize methane to CO₂ operate the methanogenetic pathway and thus the Mtr enzyme entirely in reverse.

Mtr complexes in methanoarchaea have been studied by biochemical and genetic approaches for more than thirty years (Gottschalk and Thauer, 2001). It has been known for a long time, that Mtr complexes consist of eight subunits (MtrABCDEFGH), of which the corresponding genes are organized in an operon, which is constitutively expressed (Harms *et al*., 1995). The full Mtr complex was found to have an apparent molecular mass of about 650 kDa, concluding a heterotrimeric complex (MtrABCDEFGH)_3_ (Upadhyay *et al*., 2016). This heterotrimeric complex consists of the cytosolic catalytical domains and a large membrane portion including the sodium channel. Thereby, MtrC, D and E are the largest integral membrane subunits, MtrA, B, F and G contain membrane anchors and MtrH is the soluble component in the cytoplasm (Gottschalk and Thauer, 2001).

Structural data on Mtr are currently limited to a partial cryo-EM reconstruction of the membrane subunits and an X-ray structure of the MtrA shuttle domain (Aziz *et al*., 2024; Wagner, Ermler and Shima, 2016), excluding the cytoplasmic MtrH and full-length MtrA components. While incomplete, these structures provide key insights— most notably the identification of a sodium-binding site in the MtrE subunit involving the conserved Asp187, previously implicated in sodium ion transport (Gottschalk and Thauer, 2001). A second putative Na⁺ binding site and a coenzyme M (CoM) molecule were tentatively assigned within the central MtrC cavity, though these remain to be functionally validated (Aziz *et al*., 2024). Additionally, the crystal structure of MtrA revealed a unique corrinoid-binding mode, distinct from known B12-binding proteins, utilizing a Rossmann-fold domain (Wagner, Ermler and Shima, 2016). However, the absence of a full complex structure in a catalytically competent state continues to limit mechanistic understanding of sodium-coupled methyl transfer.

While much focus has been on mechanistic studies to elucidate the mechanism of sodium translocation, relatively little attention has been paid to the regulation and integration of Mtr within the broader cellular context. In particular, the mechanisms governing Mtr regulation, as well as its potential modulation in response to environmental cues, are poorly understood.

Recent advances in ribosome profiling and quantitative proteomics have revealed an additional regulatory layer in prokaryotes, mediated by small proteins encoded by small open reading frames (sORFs). Defined as polypeptides typically shorter than 100 amino acids, these proteins are frequently omitted from genome annotations due to size thresholds, yet have emerged as widespread and functionally diverse regulators across bacteria and archaea. Expression profiling showed that many small proteins accumulate under specific growth conditions or are induced by stress arguing for a role in regulation (Burton, Zeinert and Storz, 2024). However, information regarding a specific physiological role of verified small proteins is frequently lacking for the majority of confirmed small proteins. For *M. mazei,* we recently reported on 314 previously non-annotated small ORF encoded small proteins, several of them regulated in response to nitrogen stress (Tufail *et al*., 2024). Although their precise roles remain to be established, the observed condition-dependent expression patterns raise the possibility that some may modulate key metabolic processes, including Mtr complex function or responses to environmental stressors.

In this study, we present the single-particle cryo-electron microscopy (cryo-EM) structure of the complete Mtr complex from *Methanosarcina mazei*, obtained at a resolution of 2.1 Å in C_1_. Our results detail the overall architecture of the complex, including tightly bound archaeal phospholipids and interaction of the MtrH methyltransferase subunits. Further, we identify the binding of a small protein, termed MtrI, to the cytoplasmic domain of MtrA, a previously unknown interactor conserved within the order Methanosarcinales. This binding is redox-dependent and occurs in response to cellular oxygen exposure, suggesting a potential role for MtrI in a protective mechanism against oxygen, supporting the adaptation of Methanosarcinales to microoxic environments in their natural habitats.

## Results

### The Mtr complex forms a distinct cloverleaf-shaped architecture

To achieve gentle purification of the entire Mtr complex of *M. mazei*, a plasmid-borne variant of MtrE carrying a C-terminal TwinStrep-tag (TS-tag) was introduced into wild-type *M. mazei*, facilitating one-step affinity purification. This was achieved by employing an optimized version of the Methanosarcina shuttle plasmid harboring the tagged MtrE subunit under the robust constitutive McrB promoter.

Membrane fractions obtained from methanol-grown cells harvested during the exponential phase were solubilised using either n-dodecyl β-D-maltoside (DDM) or lauryl maltose neopentyl glycol (LMNG). Following solubilization, Strep-Tactin affinity purification of the tagged MtrE successfully yielded a homogenous sample of the complete MtrA-H complex. The integrity and purity of the purified complex were analyzed by SDS-PAGE, western blotting, size-exclusion chromatography, mass photometry, and tandem mass spectrometry analysis of tryptic peptides (Supp. Fig. 1, a-e). Mass photometry determined the molecular mass to be approximately 763 kDa, indicative of the full trimeric complex embedded within a detergent micelle (supp. Fig. 1, c).

To unveil the molecular architecture of the Mtr complex, cryo-electron microscopy (cryo-EM) single-particle analysis was performed on the LMNG-solubilised sample. This analysis yielded an asymmetric reconstruction of the entire complex at a nominal resolution of 2.1 Å (Fig. 1, Supp. Fig. 2, Supp. Data Table 1). The Mtr core complex consists of a trimer of a hetero-nonameric protomers, each composed of three multi-spanning subunits (MtrCDE), four single-spanning subunits (MtrABFG), and a dimeric cytoplasmic methyltransferase MtrH. The trimerization interface is formed by the single-spanning subunits MtrABFG, creating a dodecameric core. Notably, these subunits possess an unusual length, extending twice the thickness of the membrane, forming a central stalk that protrudes towards the cytoplasm. This stalk interacts with three pseudo-symmetric heterotrimers formed by the multi-spanning subunits (MtrCDE), resulting in a three-fold symmetric, cloverleaf-shaped architecture.

**Fig. 1.**
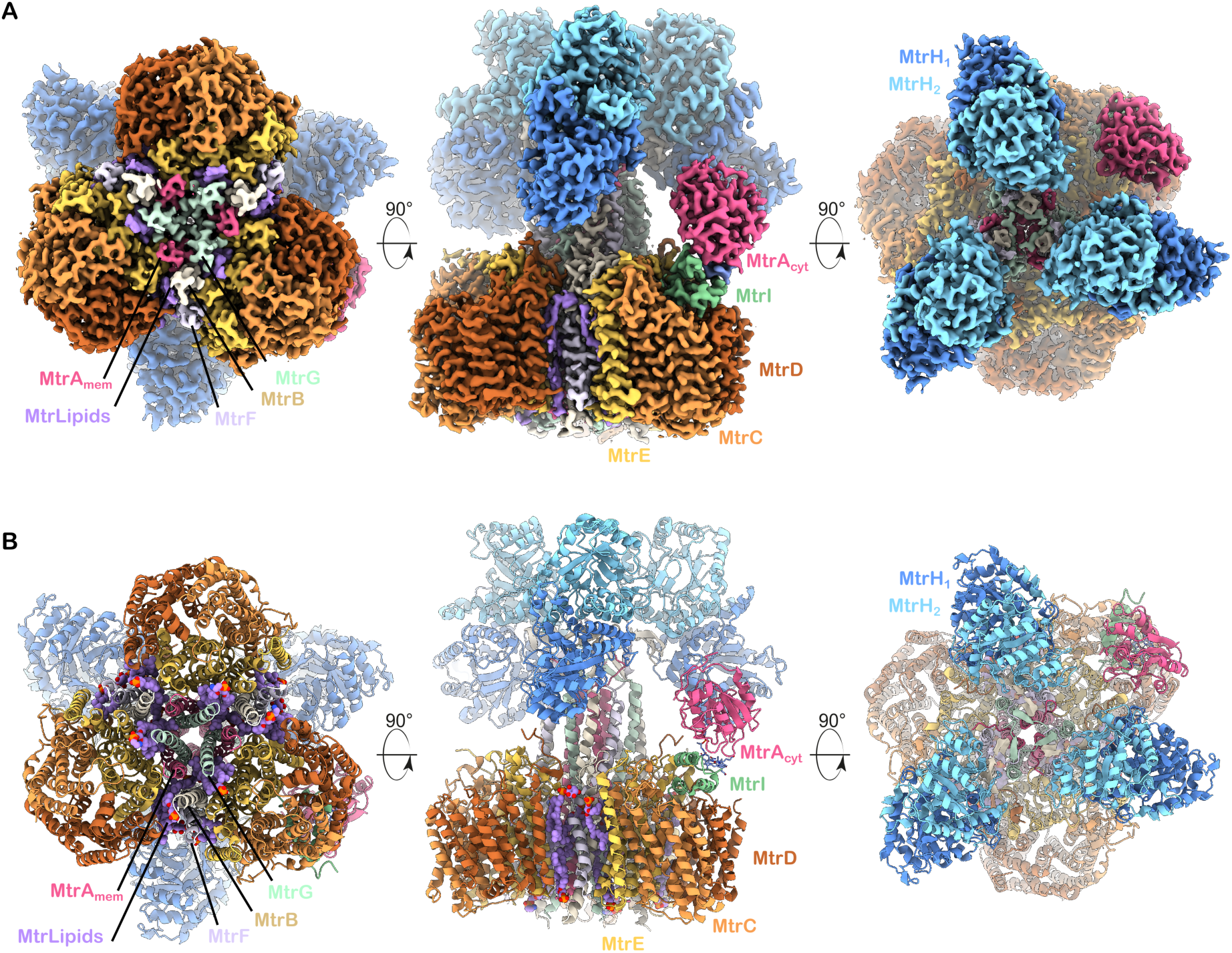
Overall architecture of the *M. mazei* Mtr complex. A, Segmented cryo-EM map of the the composite Mtr map. B. Corresponding protein model of the Mtr complex

One of the stalk subunits, MtrA, features a cytoplasmic corrinoid-binding domain, MtrA_cyt_, connected via a long, flexible linker. This domain harbours the 5-hydroxybenzimidazolylcobamide cofactor (Factor III), that serves as mobile carrier element in the methyltransferase reaction. Our cryo-EM analysis revealed that only a subset of particles exhibited a well-resolved MtrA_cyt_ domain bound to a single MtrCDE subcomplex, resulting in an overall asymmetry of the complex. Furthermore, regions encompassing MtrH and MtrA_cyt_ are highly flexible, llimiting the local resolution. To improve resolution in these flexible regions, we performed masked 3D-classification, 3D variability analysis and local refinements, yielding focused maps with resolutions of 2.5 Å for MtrA_cyt_ and 3.2 Å for MtrH. Combining these maps enabled the generation of a composite map encompassing the full complex, which in turn allowed us to obtain a complete atomic model of the asymmetric Mtr complex.

### The central stalk forms a tight hydrophobic seal that prevents ion leakage

The cytoplasmic section of the stalk (Fig. 2 A(i), B(i)), features three straight helices from chain MtrA at its core, spanning residues A:177–210. Surrounding these, the helices from chains F, B, and G are arranged in a clockwise orientation (viewed from the membrane towards the cytosol) relative to chain A, aligning almost parallel to it. Hydrophobic interactions mediate this organization, resulting in a tetrameric assembly (ABFG) on each side of the trimeric core, forming a trimer of tetramers (Fig. 2). This arrangement creates a vestibule connected to the cytoplasmic solvent, as indicated by resolved water molecules in our structure. This vestibule is closed at the border to the membrane plane by three phenylalanines (Phe210) from MtrA. In the membrane plane, the three MtrA helices bend outwards and are replaced as the core of the stalk by the MtrG helices (G44-G66), which form a three-helix bundle along the threefold axis. This bundle, characterised by conserved hydrophobic residues across methanogenic species, together with Phe210 from MtrA, acts as a tight hydrophobic seal that prevents ion leakage during conformational changes within the Mtr complex. Notably, helix B is discontinuous, featuring a cytosol-protruding loop spanning residues Pro56 to Thr70, which interacts directly with the membrane-spanning subunit MtrE. Additionally, helices F and B wrap around and cross each other between residues MtrF:42–53 and MtrB:71–80, causing a pronounced tilt of MtrF away from the threefold axis. This disruption breaks local tetrameric arrangement within the transmembrane region. Within the resulting gap, a minimum of five well-defined archaeal ether lipids are bound per trimer. The density quality allowed us to model a total of 15 lipids, though additional lipids may be present. These lipids are deeply embedded in the complex, acting as integral non-protein structural components of the Mtr complex, bridging and stabilising interactions between membrane-spanning subunits. Guided by our well-resolved cryo-EM map, the surrounding local environments, and considering the most abundant lipid species in *Methanosarcina* (Sprott et al., 1994), we modelled these embedded lipids as either archaeol or 2-hydroxy-archaeol (2,3-di-O-phytanyl-sn-glycerol) bearing the polar headgroups phosphatidylethanolamine and monophosphate or phosphatidylinositol and phosphatidylserine.

**Fig. 2.**
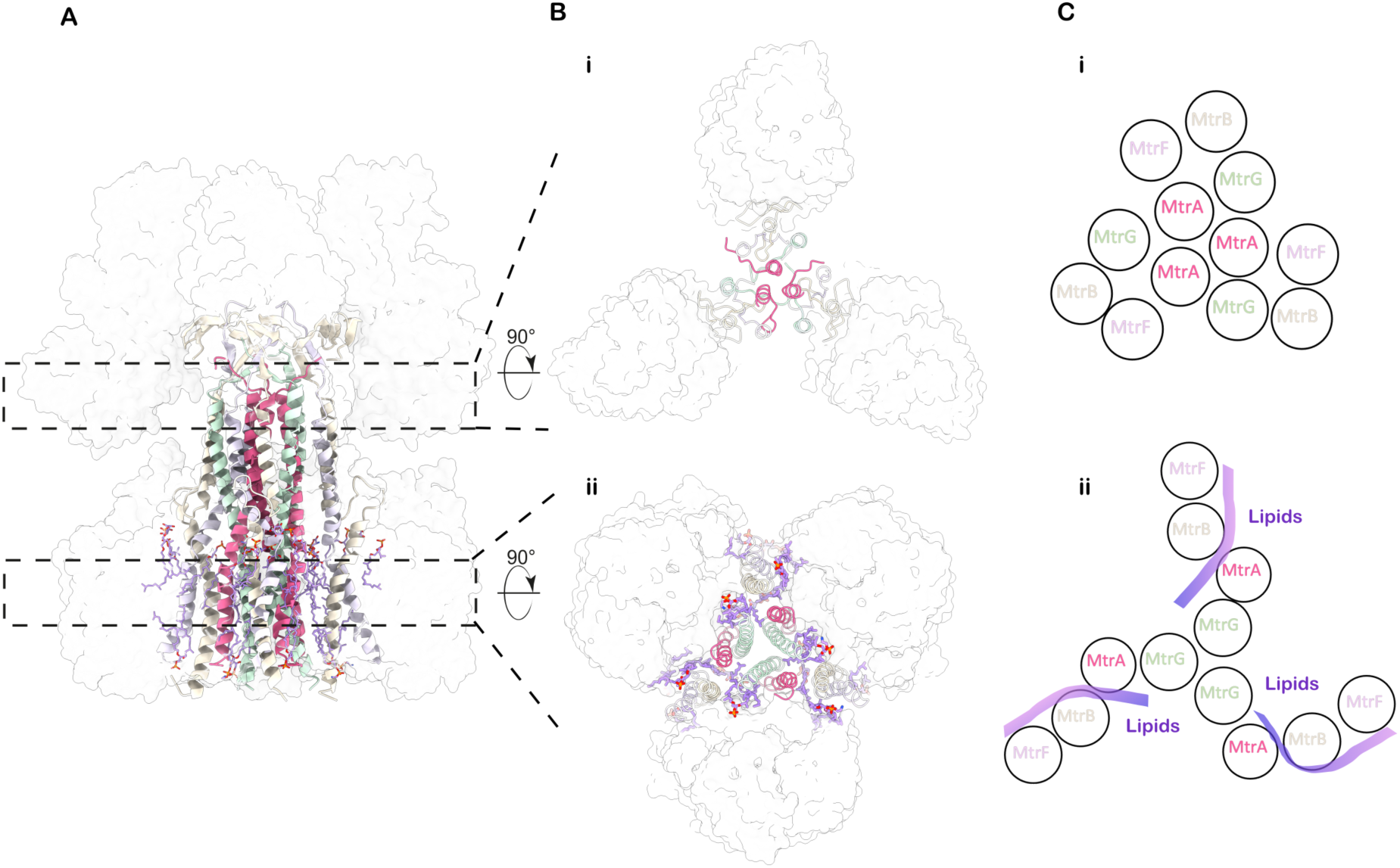
Central stalk. **A**, Side view of the central stalk. **B**, **i**) Cross-section of the stalk in the cytosolic region, **ii**) Cross-section of the stalk in the membrane region. **A**, **B**, MtrH and MtrCDE subunits are shown as white, transparent, surfaces. MtrBFG- and membrane-located MtrA-helices are shown as cartoon. Archaeal phospholipids are shown as sticks. **C**, **i**) schematic representation of subunit arrangement in the cytosolic region, **ii**) schematic representation of subunit arrangement in the membrane region

### The dimeric methyltransferase MtrH

Like other related methyltransferase systems, MtrH exists as a homodimer of two MtrH monomers, each adopting a triose-phosphate isomerase (TIM) barrel fold. The structure reveals that three MtrH dimers are bound, resulting in a total of six MtrH copies within in the complex. Similarly to its closest known structural relative MtgA (Badmann and Groll, 2020) the TIM-barrel is formed by eight parallel ß-strands in its center, enclosed and connected by helices. The MtrH monomers dimerise via their C-terminal helices. The resulting MtrH dimer associates with the end of a three-stranded β-sheet formed by the N-terminal region of MtrB (residues 1–23; Fig. 3A, B). This ensures that MtrH consistently binds with the same orientation, positioning the TIM-barrel of the membrane-proximal MtrH (MtrH_p_) to be opened in a clockwise way, as viewed from the membrane towards the cytosol. Such an orientation facilitates binding of the cytoplasmic domain of the methyl-group carrying MtrA. As the active sites of the MtrH dimer face in opposite directions, with only one oriented toward MtrA, it suggests that only the membrane-proximal MtrH_p_ is functionally relevant.

**Fig. 3.**
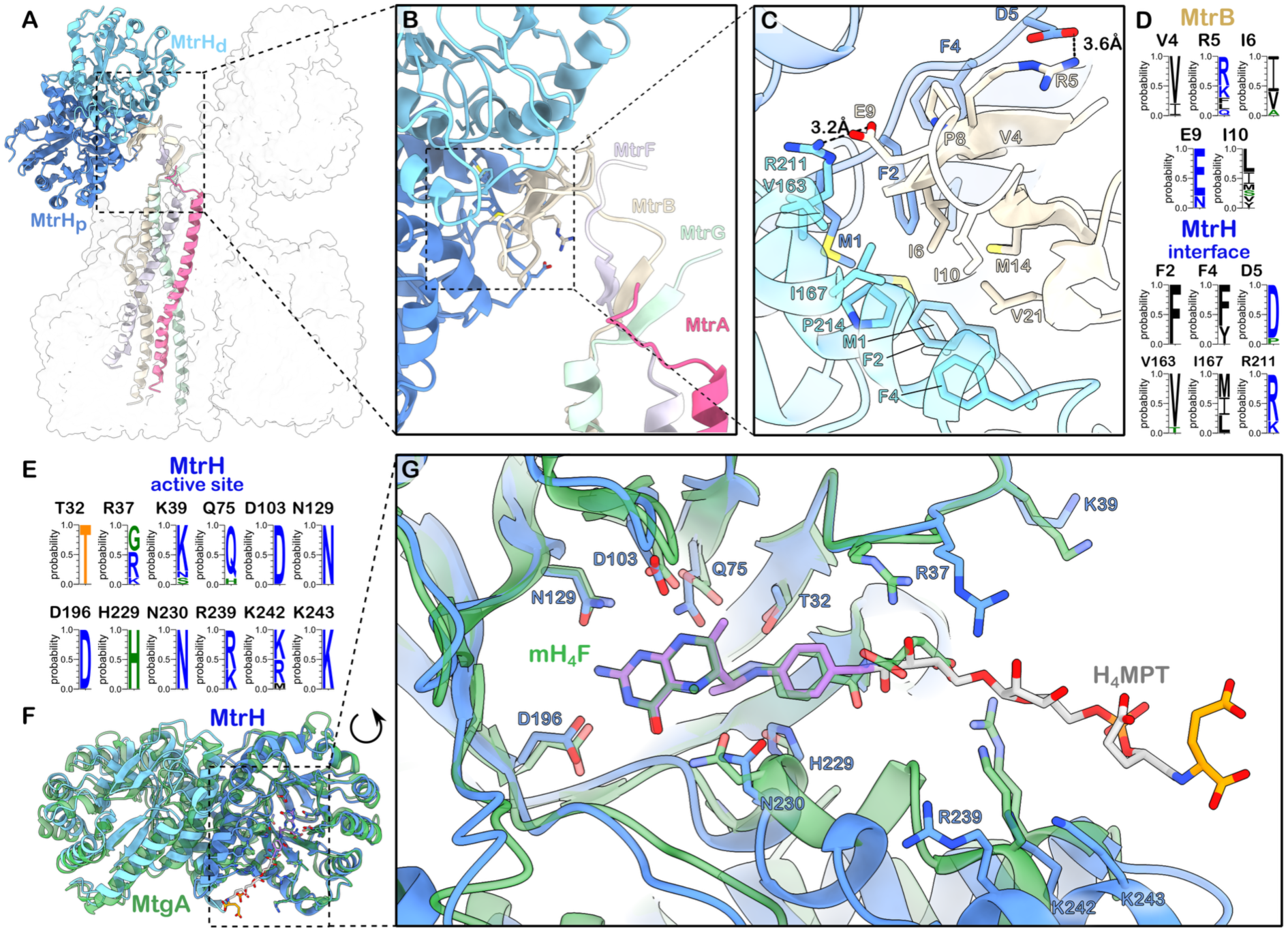
Structure and interactions of the dimeric methyltransferase MtrH. **A**, Overview of MtrH positioning in the Mtr complex. Cartoon representation of dimeric MtrH bound to one protomer of the stalk-forming MtrABFG helix-bundle. The rest of the complex is depicted as a white, semi-transparent surface. **B**, Close-up view of the extended ß-sheet formed by the N-termini of MtrBFG and the MtrH-stalk interface created by the MtrB sheet. **C**, MtrH-MtrB interface. Residues involved in hydrophobic and charged interactions are labeled and shown as sticks. **D**, Conservation of MtrB-MtrH interacting residues across Methanoarchaea. **E**, Conservation of substrate-binding residues within the MtrH active site across Methanoarchaea. **F**, Superposition of MtrH (light and dark blue) and MtgA (PDB: 6SJN, green). **G**, Close-up of the superposition of the active site of MtrH and MtgA. Residues involved and potentially involved in substrate-binding as well as Methyltetrahydrofolate (mH_4_F) and H_4_MPT are shown as sticks. The density of mH_4_F (green) bound to MtgA was used to model the binding of H_4_MPT (purple, grey, orange).

The interaction between the strands of MtrB and MtrH is mainly mediated by small hydrophobic residues that form a shape complementary to the hydrophobic binding pocket between the MtrH-dimerisation interface. In addition, it is stabilized via salt-bridges between Arg5_MtrB_ and Asp5_MtrH_ as well as Glu9_MtrB_ and Arg211_MtrH_ (Fig3C, D). This rather weak interface also explains the often occurring loss of the methyltransferase in native purification procedures (Upadhyay *et al*., 2016).

To better understand Tetrahydrosarcinapterin (H₄SPT) binding by MtrH, we superposed our MtrH structure with the structure of the homologous methyltransferase MtgA in complex with Methyltetrahydrofolate (mH_4_F) (Badmann and Groll, 2020, PDB: 6SJN). Although the ligand is not present in the structure, the superposition reveals a highly conserved arrangement of residues within the pterin-binding site, suggesting a conserved mode of pterin-moiety recognition. Residues Asp103, Asn129, Asp196, and Asn230 in MtrH are conserved and correspond to key residues in MtgA, suggesting they play identical roles in both methyltransferases. Additionally, Thr32, Gln75, and His229, which also line the pterin-binding site, are not strictly conserved but preserve properties of their counterparts in MtgA and are therefore likely to fulfill analogous functions. This conservation of the binding environment enabled us to model H₄SPT into the substrate binding-pocket of MtrH by superposing the mH_4_F ligand from MtgA. Further, comparative sequence analysis across methanogenic archaea identified an additional conserved feature: a positively charged patch formed by residues Arg239, Lys242, Lys243 at the exit site of the enzymatic cavity. This conserved patch suggests a potential additional binding interface accommodating the polar and negatively charged ribosyl and gamma/alpha-glutamyl tail of H_4_MPT/SPT. However, confirmation of the exact substrate-binding mode will require an experimental substrate-bound structure of MtrH.

### The CED Trimer and the sodium ion binding site

Three integral membrane MtrCDE trimers are attached symmetrically around the central MtrABFG stalk (Fig. 4A). The MtrCDE trimer interacts with all components of the central stalk, except for the trimeric core-forming MtrG, and are stabilized by five lipids. Within the MtrCDE trimer only MtrE directly interacts with the stalk subunits, while MtrC and MtrD are positioned on the exterior of the protein complex. Notably, transmembrane helices TM7 and TM8 of MtrD interact specifically with 2-hydroxy-archaeolphosphatidylinositol and archaeolphosphatidylethanolamine lipids.

**Fig. 4.**
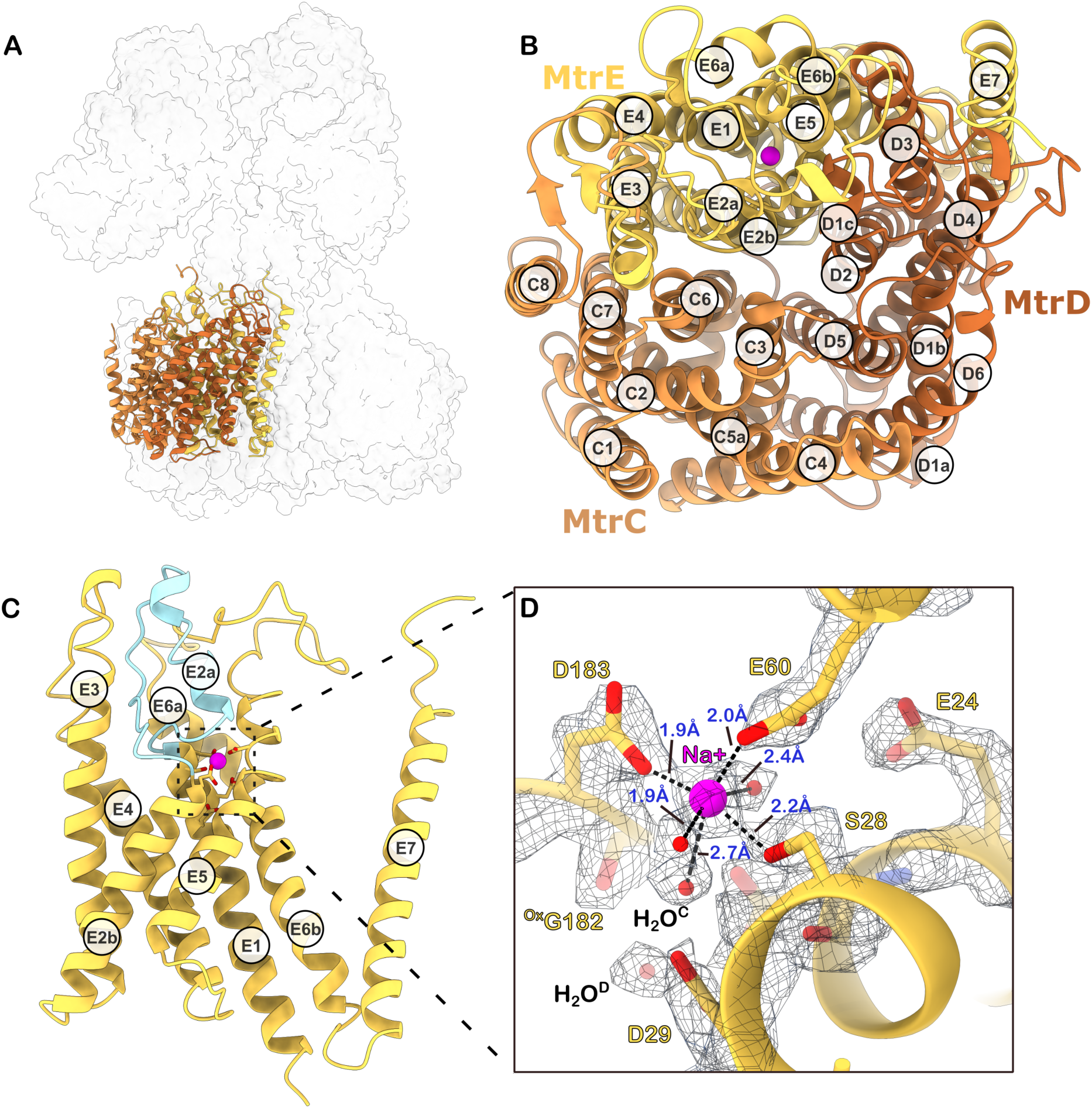
The CED membrane globe and sodium ion binding site. **A**, Overview of a single CED trimer in Mtr complex. **B,** CED trimer (cartoon) with helices labelled. Top-down view from the cytosolic side towards the membrane **C,** Side view of MtrE (cartoon). Membrane-embedded loop-helix-loop structure highlighted in light blue. **D,** Close-up view of the sodium ion binding site within MtrE. Selected residues shown as sticks together with their cryo-EM density (threshold: 0.116). D183, E60 and S28 together with three water molecules form an octahedral coordination complex around a sodium ion. A previously proposed second sodium ion has been reassigned as a water H_2_O^D^.

MtrC, MtrD, and MtrE are each composed of six transmembrane helices organized into a pseudo-twofold symmetrical helical bundle (Fig. 4B). This internal symmetry allows TM1-3 to structurally superimpose onto TM4-6. Structural comparisons using DALI reveal that these proteins possess a rare fold with no close structural analogs among other membrane proteins. However, modest similarities exist: MtrE shows resemblance to the heme transporter HmuUV, the vitamin B12 import system permease protein BtuC, and, intriguingly, the sodium-translocating RnfE. Despite low sequence identity, the resemblance to RnfE raises the possibility that this fold may have evolved to mediate sodium translocation under energy-limited conditions.

Interestingly, despite the shared overall architecture of MtrC, MtrD and MtrE, the sequence identity between these proteins is relatively low (MtrC–MtrE: 21.36%; MtrD– MtrE: 21.43%; MtrC–MtrD: 25%). This indicates these subunits likely descended from a common ancestor but diverged along distinct evolutionary paths, leading to specialized functions within the complex.

Previously (Gottschalk and Thauer, 2001), sequence conservation analysis and the hallmark feature of high charge suggested that a signature loop in MtrE was a cytoplasmic loop containing a zinc-binding area. Contrary to this, our structural analysis reveals that this region in MtrE is actually a membrane-embedded element with a loop-helix-loop structure (Fig. 4C). The previously proposed highly conserved zinc-binding residues Asp25, Asp29, and Glu60, along with the suggested sodium ionbinding site residues Asp183 and Ser28, are in close proximity, forming a hydrophilic, charged pocket within MtrE. This pocket likely forms the sodium ion binding site of the Mtr complex. Our structure shows density consistent with a sodium ion octahedrally coordinated by conserved residues (Fig. 4D). This finding is in agreement with a previously reported Mtr structure from *Methanothermobacter marburgensis* (Aziz *et al*., 2024, PDB: 8Q3V), highlighting the conservation of this binding sites in all methanogens.

### The small protein MtrI bridges MtrA_cyt_ and the MtrCDE subcomplex

Unexpectedly, in our structure, the corrinoid-binding domain of MtrA does not directly interact with the integral membrane MtrCDE subcomplex. Instead, our cryo-EM map indicates that this interaction is bridged by a previously unidentified small helical protein (Fig. 5A). We employed ModelAngelo (v.1.0) (Jamali *et al*., 2024) to automatically build a model without a sequence input. Using BLASTP with the output-sequence from ModelAngelo, the helical protein was identified as MM_2401, a small protein consisting of only 68 amino acids (Supp. Fig. 3d). The respective gene is not part of the highly conserved *mtr* operon but represents a stand-alone gene and has so far evaded detection and attention. We named this protein MtrI.

**Fig. 5.**
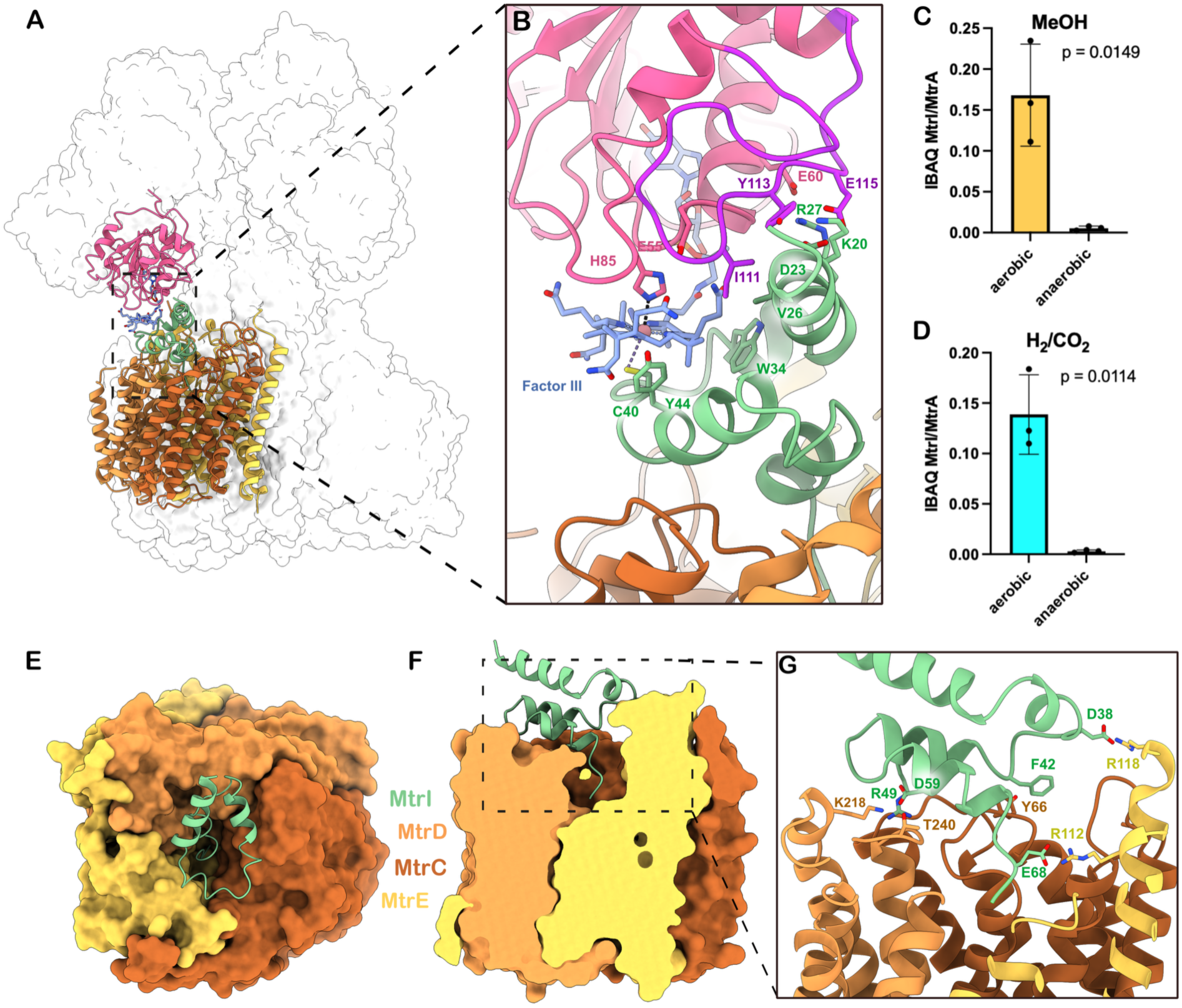
MtrI binding to MtrCDE and MtrA is oxygen-dependent. **A**, Overview of MtrI (green) location in Mtr. MtrI is sandwiched by MtrA_cyt_ (pink) and MtrCDE (brown, yellow, orange) **B**, Close-up of MtrA-MtrI interaction. Interacting residues and Factor III shown as sticks and labeled. Unstructured insertion domain of MtrA is colored purple. **C**, Ratio of iBAQ values of MtrI and MtrA of Mtr complex purified under aerobic and anaerobic conditions from methanol grown cells. **D**, Ratio of iBAQ values of MtrI and MtrA of Mtr complex purified under aerobic and anaerobic conditions from H_2_/CO_2_ grown cells. **E**, Top view of MtrI binding to the MtrCDE membrane globe (surface representation). **F**, Side view of MtrI binding to the MtrCDE membrane globe (surface representation, clipped). **G**, Close-up of MtrI-MtrCDE interaction. Interacting residues are shown as sticks and labeled.

MtrI consists of four helices (Fig. 5B, Supp. Fig. 3e) that are lying atop the cavity of the MtrCDE subcomplex (Fig. 5E, F) stabilized by a series of charged and aromatic interactions (Fig. 5D): Asp38 interacts with Arg118 of MtrE, Phe42 forms aromatic interactions with Tyr66 of MtrC, Arg49 binds Lys224 of MtrD via a bridging water and Asp59 interacts with Thr240 of MtrD. Crucially, the C-terminal residues of MtrI penetrate into the vestibule formed by the three MtrCDE subunits, where it is anchored by an interaction between the C-terminal Glu67 of MtrI and Arg112 of MtrE. This arrangement effectively blocks the central cavity of MtrCDE (Fig. 5F). Within the other two MtrCDE subcomplexes of the Mtr, we do not observe any MtrI. A stronger density next to Arg112 could be explained in accordance with another study (Aziz *et al*., 2024) by a mixture of two partially occupied CoM sites. The presence of MtrI in the cavity, along with its interaction with Arg112, therefore likely occludes the CoM binding site and blocks the sodium channel.

Superimposing MtrCDE not containing MtrI with the site of the trimer showing the MtrCDE–MtrI complex reveals no major structural differences. One notable exception is a slight rearrangement of the cytosolic loop in MtrC (residues 65–71), which includes Tyr66, to accommodate MtrI binding. This loop also appears more rigid in the cryo-EM density of the MtrCDE–MtrI complex.

Interestingly, MtrI features an N-terminal Zinc-ribbon domain comprising 15 residues, including 4 cysteines that form a zinc-binding motif (Supp. Fig. 3e). However, in our cryo-EM map, the N-terminal region containing the Zinc-ribbon is highly flexible and poorly defined. Thus, we recombinantly expressed N-terminally TwinStrep-SUMO tagged MtrI in *Escherichia coli*, purified the small protein and successfully demonstrated its Zinc-binding properties using ICP-MS (Supp. Fig. 3f-i). Like most zinc-ribbons, this one likely serves a structural role, although its exact function remains unknown.

To further validate binding of MtrI to the Mtr complex we performed a reciprocal pulldown experiment by expressing an N-terminal His_6_-tagged variant of MtrI from a modified *M. mazei* shuttle plasmid in *M. mazei* (Supp. Fig 3b). Ni-NTA purification of these strains resulted in strong enrichment of Mtr subunits compared to the empty vector control (Supp. Fig. 4a, b).

MtrA_cyt_ of *M. mazei* shows a high degree of structural similarity to the previously reported structure from *Methanothermus fervidus* with an RMSD of 0.599. As noted earlier, MtrA exhibits a distinctive binding mode of the corrinoid cofactor Factor III (FIII), setting it apart from other enzymatic systems (Wagner, Ermler and Shima, 2016). A key feature of the interaction between MtrA and MtrI is the direct coordination of the cobalt atom of FIII by Cys40 of MtrI, which serves as the upper axial ligand (2.67 Å) and His85 of MtrA, which acts as the lower axial ligand (2.68 Å). Additional contacts are formed between Tyr44 and Trp34 of MtrI and the porphyrin ring of FIII (Fig. 5B). These interactions are further stabilized by a network of polar contacts involving Lys20, Asp23 and Arg27 of MtrI and the MtrA residues Tyr113 and Glu115 of the characteristic unstructured insertion domain of MtrA (residues 98-120), as well as Glu60.

### MtrI binding is redox dependent and is triggered by oxygen exposure

The unexpected presence of MtrI in our cryo-EM structure prompted us to investigate the significance of its binding on the function of the Mtr complex. If MtrI were essential for the core function of the Mtr complex and directly involved in sodium-translocation, one might expect it to be universally conserved across all methanogens. However, BLAST analysis revealed that MtrI is uniquely present in the order Methanosarcinales.

When comparing Methanosarcinales with other methanogenic orders, two major differences stand out: (i) Methanosarcinales can grow on additional substrates, facilitated by the presence of methanophenazines and additional energy-conserving systems that directly affect Mtr function, which e.g. is reversed during methylotrophic growth (methanol, methylamines), and (ii) Methanosarcinales often inhabit oxygen-rich environments. To assess whether MtrI plays a function in mediating one of these physiological traits we grew *M. Mazei* containing the MtrE-TS plasmid on either methanol or H_2_/CO_2_ and subsequently purified Mtr from these cultures under both aerobic or strictly anaerobic conditions.

To quantify MtrI abundance we performed liquid chromatography-tandem mass spectrometry (LC-MS/MS) analysis of tryptic peptides derived from purified Mtr. Protein intensities and IBAQ values were calculated using MaxQuant (v.2.6.5.0) (Cox and Mann, 2008) and the MtrI:MtrA iBAQ ratio was used as a semi-quantitative metric to determine MtrI levels. For both growth conditions, H₂/CO₂ and methanol, aerobic purification of the Mtr complex consistently yielded similar levels of MtrI, which remained within the same order of magnitude (Fig. 5C, D). These aerobically purified samples appeared pink, indicating an oxidized Co(III) state. In contrast, Mtr samples purified under strictly anoxic conditions - yielding a brown-coloured sample that indicates a reduced Co(II) or Co(I) state - exhibited markedly lower levels of MtrI across both growth conditions (Fig. 5C,D). These findings suggest that MtrI association requires the oxidized Co(III) state of the corrinoid cofactor. Thus, MtrI binding to the Mtr complex appears to be redox-dependent and may be triggered within the cell upon exposure to oxygen.

## Discussion

This study presents the first structural evidence of a small protein associated with a core bioenergetic complex in Methanoarchaea. This highlights the regulatory potential of small proteins, a class of the proteome that has been historically overlooked due to methodological constraints. Furthermore, our findings substantially advance the mechanistic understanding of the Mtr complex by providing a highly resolved cryo-EM structure in which all subunits are visible, offering new insights into its assembly and organisation. In contrast to previous suggestions of partial binding, our results reveal full occupancy of an MtrH dimer in the stalk region, which clarifies the dimer’s role in the architecture of the complex. This complete structure has also identified a single sodium-binding site, critical for understanding how sodium ion pumping is integrated with methyl transfer reactions in the energy conservation processes of methanogens. Additionally, the structure implicates two possible regions were MtrA may transiently bind - one at the membrane-proximal MtrH_p_ TIM-barrel and another at the cavity formed by the MtrCDE trimer, where a potential CoM binding site has been proposed. Such a movement of MtrA could play a role in coupling the methyl transfer reaction with sodium ion pumping through conformational changes. However, despite these advancements, several critical gaps remain in our understanding of how these processes are fully coupled. To resolve this, further structural studies are needed under active turnover conditions.

A particularly exciting discovery was the identification of MtrI, a previously uncharacterized small protein encoded by an sORF, tightly associated with the Mtr complex in our preparation. This association was confirmed by reciprocal affinity purification of the Mtr complex using a His_6_-tagged variant of MtrI expressed in *M. mazei*. Our structure further suggests that MtrI may recognize oxidatively modified or inactive forms of MtrA in the Mtr complex. It is known, that the number of axial ligands that bind to cobalt in cobamide-cofactors is dependent on the redox-state of the cobalt center. In general, Co(I) is ligand-free, Co(II) harbors one axial ligand (either upper or lower) and Co(III) binds two axial ligands (upper and or lower). The presence of Cys40 from MtrI as an upper-axial ligand and His85 of MtrA as a lower-axial ligand suggests a Co(III) oxidation state. This oxidation likely results from oxygen exposure during purification, suggesting that MtrI senses an oxidized cobamide cofactor. By blocking the MtrCDE cavity, MtrI might prevent sodium leakage and inhibit MtrA-CoM turnover. The stability of the Mtr complex under these conditions is uncertain. It could be marked for repair by connectase enzymes (Fuchs *et al*., 2021), activated reductively by as-yet uncharacterized ATP-dependent RaCo enzymes (Huening, Jiang and Krzycki, 2020), or targeted for degradation. While the exact function of MtrI remains unclear, its association with Methanosarcinales’ adaptation to oxygen exposure is particularly intriguing.

Unlike Class I methanogens, Methanosarcinales exhibit greater resilience to microoxic conditions and inhabit diverse environments, such as wetlands, peat bogs, and oxygenated sandy sediments, where they frequently encounter oxygen exposure. The exclusive occurrence of MtrI in Methanosarcinales suggests that it evolved as an adaptation to oxygen-exposed environments. *Methanosarcina* are also known to employ mechanisms such as antioxidant enzymes, including catalase and superoxide reductase, as well as a high-affinity terminal oxidase, cytochrome bd (CydAB), to contend with oxygen and oxidative stress. Changes in thiol-molecule and polyphosphate (polyP) content, along with the development of biofilms, further enhance their ability to withstand oxidative conditions. Further studies are essential to clarify MtrI’s precise role and determine whether it represents a key factor in oxygen resistance, potentially as part of a broader protective mechanism within Methanosarcinales. Besides our finding, there is further recent evidence that small proteins can modulate the activity of the Mtr complex (Habenicht *et al*., 2025).

Overall, this study has an impact that extends beyond our understanding of Mtr: it establishes the previously unrecognized role of small proteins in regulating core bioenergetic complexes in methanogenic archaea. By revealing MtrI to be a redox-responsive modulator of the Mtr complex, the study sets a precedent for small proteins as key regulatory elements in the methanogenic metabolism.

## Material and Methods

### Chemicals

Unless specified otherwise, chemicals were acquired from Sigma-Aldrich, Carl Roth, Roche and Serva GmbH

### Strains and plasmids

MtrE was expressed with a C-terminal TwinStrep tag (TS-tag) in *Methanosarcina mazei* DSM #3647 using plasmid pRS1743. pRS1743 was generated via Gibson Assembly and consists of the backbone pRS1595 (a shuttle vector for *M. mazei* and *Escherichia coli*; Thomsen and Schmitz, 2022), the constitutive promotor *pmcr*B including a ribosome binding site (RBS) as control element for the C-terminally TS-tagged *mtr*E gene (*MM_1547*) (Supp. Fig. 3a). In detail, the backbone pRS1595 was restricted with NotI and BamHI (New England Biolabs, Ipswich, USA) to obtain a linear fragment. Primers for the PCR-products (*pmcr*B, *mtr*E and TS-tag) were generated using NEBuilder (New England Biolabs, Ipswich, USA) and commercially synthesized by Eurofins Genomics (Ebersberg, Germany). The promotor *pmcr*B was PCR-amplified using the template pRS893 and primers Gibson_Promotor_for (5’agggccctaggtaccatatgGAGCTCTGTCCCTAAAAATTAAATTTTC3’) and Gibson_Promotor_MtrE_rev (5’gtggttccatGTTTAATTTCCTCCTTAATTTATTAAAATCAC3’); the *mtr*E -gene was amplified using chromosomal *M. mazei* DSM #3647 DNA as template and primers Gibson_MtrE_for (5’gaaattaaacATGGAACCACTCATAGGCATG3’) and Gibson_MtrE_rev (5’accacgctgaTGCGGAAGCCTCCTCAGC3’); the TS-tag was PCR-amplified using template plasmid JS004 (a vector based on pET28a) and primers Gibson_twinstrep_MtrE_for (5’ggcttccgcaTCAGCGTGGTCGCATCCC3’) and Gibson_twinstrep_rev (5’actagtaacgttaagcttgc**AAAAAAA**TTACTTTTCAAATTGAGGATGGGACC3’). The TS-tag reverse-primer contained additional A_7_ that function as a transcriptional terminator in *M. mazei* (T_7_ on mRNA level). The Gibson Assembly reaction was performed by using the Gibson Assembly® Master Mix (New England Biolabs, Ipswich, USA), 30 fmol restricted pRS1595, 90 fmol *mtr*E -PCR product and 210 fmol of *pmcr*B - and TS-tag-PCR-products each for 60 min at 50 °C. The assembly was then transformed into *E. coli* DH5α λpir following the method of Inoue (Inoue, Nojima and Okayama, 1990). The resulting plasmid pRS1743, which contained a C-terminally TS-tagged *mtr*E under control of the *pmcr*B promotor, was then transformed into *M. mazei*\* cells and a single clone was isolated as described in Ehlers *et al*., 2005.

His_6_-MtrI was expressed in *M. mazei* using plasmid pRS2145 (Supp. Fig. 3b). pRS2145 was generated by PCR-amplification of *His_6_-MtrI* from template pRS2139 (Hüttermann and Schmitz, 2024) using primers His_MM_2401_NdeI_for (5’TTTCATATGCACCATCACCATCATCACATGC3’) and His_MM_2401_NheI_rev (5’TTTGCTAGCTCAGTCCTCGACGTCAAGAG3’). The PCR-product and pRS1807 were restricted using NdeI and NheI (New England Biolabs, Ipswich, USA), ligated and transformed into *E. coli* Pir1 cells (Thermo Fisher, Waltham, USA). The resulting plasmid pRS2145 was transformed into *M. mazei*\* cells by liposome-mediated transformation with sucrose according to Ehlers *et al*., 2005.

MtrI (MM_2401) was expressed with an N-terminal TwinStrep-SUMO-tag (TS-SUMO) in *Escherichia coli* strain BL21 (DE3) from plasmid pEZ13. The plasmid was generated via Golden Gate Assembly from backbone JS004 (a vector based on pET28a). Coding sequence *mm_2401* was amplified from the *Methanosarcina mazei* DSM #3647 genome with forward (5’atgcaggtctcaatgaAATGTGAAGCATGTG3’) and reverse (5’tacgtggtctcttcgaTCAGTCCTCGACGTC3’) primers, and the SUMO coding sequence was amplified from plasmid pET28-His6-SUMO with forward (5’atgcaggtctcacatgTCGGACTCAGAAGTC3’) and reverse (5’atgcaggtctcatcatTCCACCAATCTGTTCTCTG3’) primers, containing matching directional overhangs produced upon BsaI cleavage. The plasmid was assembled using BsaI-HF®v2 and T4 DNA Ligase (NEB).

### Cultivation and harvest

*Methanosarcina mazei* Gö1 was grown at 37 °C, in closed, anaerobic growth tubes in volumes ranging from small volumes (e.g., 5 mL in Hungate tubes) to larger volumes of up to 1 L in 2 L Duran bottles. The medium used was similar to DSMZ120 but buffered with 20 mM PIPES (pH 7.0) instead of NaHCO_3_ and for cultures containing plasmids puromycin was added at a final concentration of 2-5 mg/L (Alomone Labs). Media were prepared aerobically and anaerobised after autoclaving by repeatedly cycling the gas-phase with N_2_. Complete anaerobisation was achieved by addition of 40 mg/L of Na_2_S followed by visually monitoring the reduction of resazurin. Vitamins, minerals and antibiotics were added after autoclaving. Growth substrates used were either 250 mM methanol or an 80%/20% H_2_/CO_2_ atmosphere. When growing on H_2_/CO_2_, cultivation was performed in an orbital shaking incubator at 80-100 rpm. In general, growth was monitored by determining the optical density of the cultures at 600 nm (OD600). Cells were harvested during late log-phase at an OD600 of 0.7-1 at 10 000 x g at 4 °C. For anaerobic harvest centrifugation bottles and lids were pre-incubated for at least 16 h under strict anaerobic conditions inside of a vinyl anaerobic chamber with 95% N_2_ and 5% H_2_ (Coy Laboratory Products). Cultures were shuttled into the chamber and successively transferred into the centrifugation bottles. Tightly sealed bottles were moved outside the tent for centrifugation. Resazurin in the growth media served as an indicator to confirm maintenance of anaerobic conditions throughout the process. Harvested cells were stored at -80 °C.

### Purification of the Mtr complex via MtrE-TwinStrep Affinity Tag for subsequent Cryo-EM

Cells were suspended in lysis buffer [Buffer A (50 mM MOPS/NaOH pH 7.0, 10 mM MgCl_2_, 150 mM NaCl) and 0.1 mg/ml DnaseI, cOmplete™ Protease Inhibitor Cocktail] in a volume ratio of 1 g cells per 10 ml of buffer. Cells were lysed by French press (1 000 - 1 200 psi) up to 3 times at 4°C. The lysate was initially centrifuged at 20 000 x g for 30 min at 4 °C to remove cell-debris. For preparation of the membrane the supernatant was further centrifuged at 100 000 x g for 1.5 h at 4 °C. The membrane pellet was solubilised in solubilization buffer containing lysate buffer together with either 2.5 % w/v n-Dodecyl β-D-Maltosid (DDM) or 1.5 % w/v Lauryl Maltose Neopentyl Glycol (LMNG). The volume was chosen to yield a 1:2 w/w ratio detergent/membrane-pellet. Solubilization was performed for 12-16 h at 4 °C on a slowly rotating roller mixer. Non-solubilised membrane was removed by a second round of ultracentrifugation (100 000 x g for 1.5 h at 4 °C). The complex was further purified using a gravity column containing equilibrated Strep-Tactin Sepharose resin (IBA Lifesciences). The supernatant was incubated with Strep-Tactin Sepharose resin (IBA Lifesciences) for 1-2 h before its transfer to a gravity column. For the purification using LMNG, the resin was washed with Buffer A and Mtr complex was eluted with Elution Buffer (Buffer A plus 2.5 mM desthiobiotin). For the purification using DDM, 0.05% w/v DDM was added to Buffer A and Elution Buffer. The protein complex was concentrated with a 100-kDa Amicon Ultra centrifugal filter (Merck Millipore). Protein concentration was determined by Bradford assay, using a calibration curve created with bovine serum albumin. Non-heated samples were analysed by 15% SDS-gels to determine protein purity and quality. Western-blot with a horseradish peroxidase (HRP)-coupled anti-TwinStrep-tag antibody (IBA Lifesciences) was performed to visualize and confirm the presence of TS-tagged MtrE.

### TwinStrep-SUMO-MtrI expression in *E. coli*

For anaerobic protein production of TS-SUMO-MtrI, pEZ13 was transformed into *E. coli* BL21 (DE3) by the heat-shock method. Cells were cultivated in Lysogeny broth (LB). *E. coli* cultures were transferred into glass bottles upon reaching OD600 of 0.6-0.8, supplemented with 25 mM sodium fumarate, 0.5 % (w/v) glucose, 1 mM L-cysteine, 1 mM ferric ammonium citrate, 50 mM MOPS, pH 7.4, and 0.001 % (w/v) resazurin. Bottles were sealed with rubber stoppers and the gas atmosphere exchanged with nitrogen. Protein expression was induced by the addition of 0.5 mM IPTG, followed by overnight cultivation at 20 °C. Protein was purified in an anaerobic chamber filled with 95 % N2, 5 % H2, with anaerobised buffer. Cells were harvested by centrifugation and lysed using a Microfluidizer™ (Microfluidics™) in buffer (50 mM MOPS, pH 7.0, 10 mM MgCl2, 150 mM NaCl) with added lysozyme, DNase, and 0.5 mM PMSF. Clear lysate was applied to Strep-Tactin® Sepharose (IBA Lifesciences), washed, and eluted with 2.5 mM Desthiobiotin (IBA Lifesciences). Aerobically, His6-Ulp1 was used to cleave off the TS-SUMO tag, and concomitantly removed by passing the eluate over HIS-Select® Nickel affinity gel (Merck). Protein was further purified by gel filtration using a Superdex™ 75 Increase 10/300 (cytiva), concentrated using a 3 kDa MWCO Amicon® (Merck Millipore), and analysed by Coomassie-stained SDS-PAGE. Metal content was analysed by ICP-MS.

### His_6_-MtrI expression and purification in *M. mazei*

His_6_-MtrI was expressed from plasmid pRS2145 in *M. mazei* and purified via His-tag-affinity chromatography-purification. The His_6_-MtrI-expressing cultures were grown in 1 L cultures on MeOH to an optical turbidity at 600 nm (OD600) of 0.65–0.85. From now on, each step was performed aerobically. Cells were harvested (6371 × g, 45 min, 4 °C), resuspended in 5 ml buffer A (50 mM MOPS/NaOH pH 7, 10 mM MgCl_2_, 150 mM NaCl (Chemicals Carl Roth GmbH + Co. KG, Karlsruhe, Germany) and cell disruption was performed twice using a French Pressure Cell at 4.135×106 N/m^2^ (Sim-Aminco Spectronic Instruments, Dallas, Texas) followed by centrifugation of the cell lysate for 30 min (6,000 × g, 4 °C). The supernatant cell extract was solubilised using 1 % LMNG (Thermo Fisher, Waltham, USA) at 4°C overnight, followed by a centrifugation step to remove insoluble debris (30 min, 6,000 × g, 4 °C). His-tag-affinity chromatography-purification was performed with a Ni-NTA agarose (Cube Biotech, Monheim, Germany) gravity flow column. The column-bound protein was washed with 20 and 50 mM imidazole (SERVA, Heidelberg, Deutschland) and the protein was eluted with 100 and 250 mM imidazole. The aerobic purification was also performed with *M. mazei* cells containing the empty plasmid pRS1595 as a control.

### Inductively coupled plasma mass spectrometry (ICP-MS) analysis

ICP-MS was used to measure metal-concentrations. Briefly, acid digestion of samples was performed in 11% (v/v) HNO_3_ (Suprapur, Merck), for 3 h at 80 °C. This was followed by a 5.5 x dilution with Chelex-treated water, yielding a total volume of 250 µL per sample. Internal standards, indium (2 ppb) and germanium (20 ppb) (both from VWR Merck), were added to each sample for normalization. Elemental analysis was conducted using an inductively coupled plasma mass spectrometer iCAP Q (ICP-MS, Thermo Fisher Scientific), equipped with an SC4DX autosampler (Elemental Scientific) and a MicroFlow PFA-100 nebulizer. Quantification of metals was done by comparison with serial dilutions of the ICP multi element standard solution XVI (Merck). The ICP-MS operated with a reaction cell using a helium/hydrogen gas mixture (93/7%). Data acquisition was performed in triplicate using Qtegra software v2.18 (Thermo Fisher Scientific). Blank values and quality thresholds were calculated using protein-buffer standards. The measured concentrations (initially in ppb) were converted to molar units (µM) of metal per sample for quantitative analysis.

### Determination of substrate- and O_2_-dependence of MtrI binding

Cells harboring the MtrE-TS affinity tag plasmid were grown in triplicates in 2L glass bottles either on Methanol (1 L total culture volume) or H_2_/CO_2_ (0.75 L total culture volume) to an OD600 of 0.6-1. Upon reaching the target OD, cultures were transferred to an anaerobic chamber (95% N_2_ and 5% H_2_). Inside the chamber, two culture bottles were carefully combined and split again into two equal halves. Successively, one half was taken outside for aerobic harvest and the other half was filled into anaerobised centrifuge bottles and taken outside for harvest anaerobic harvest. Aerobically harvested cells were exposed to atmospheric oxygen for 1.5 to 2h before flash-freezing in liquid N_2_. Cells were stored at -80 °C under aerobic or anaerobic conditions, respectively. To quantify MtrI bound to the Mtr complex in the presence or absence of oxygen, Mtr was purified from anaerobically harvested cells under strict anaerobic conditions and from aerobically harvested cells under aerobic conditions. Mtr complex was purified as described above but in order to simplify the procedure and reduce the variations between anaerobic and aerobic purification a few steps in the protocol were changed. To lyse the cells, instead of a French press a sonicator was used, with 3 times 3 min on/ 3 min off cycle at 50-60% power and a pulse-length of 0.5s/s. Further, solubilisation was done directly with the cleared lysate omitting both ultracentrifugation steps by adding 1.5% LMNG (final concentration) and incubating for 12-16 h.

### Protein mass spectrometry, peptide identification and quantification using MaxQuant

For the mass spectrometry of purified Mtr complex 10 µl of concentrated Mtr complex was treated with 1% sodium lauroyl sarcosinate (SLS) and 1.5 µl of TCEP, heated for 10 min at 90 °C. Subsequently, 1.5 µl of 0.4 M iodacetamide was added for modification, followed by sample cleanup using sp3-beads. Samples were successively Trypsin digested overnight, desalted with C18 columns (Macherey Nagel, Chromabond C18 WP), dryed in a Speedvac and reconstituted in 0.1 % TFA.

Peptides were analysed using liquid chromatography-mass spectrometry in an Orbitrap Exploris 480 (Thermo Fisher Scientific) coupled to an UltiMate 3000 RSLCnano system. For each sample 1-4 µl of the peptides were injected onto a C18 reverse-phase HPLC column using a 30-min gradient (0.15% formic acid to 0.15% formic acid with 35% acetonitrile). The quantity of peptides injected was consistent across different samples within each experiment: 0.06 µg for H_2_/CO_2_ samples, 0.1 µg for methanol samples and 0.1 µg for His_6_-MtrI-pulldown samples. The mass spectrometry data were analysed initially using Proteome Discoverer 1.4 (Thermo Fisher Scientific) against the proteome of *Methanosarcina mazei* (DSM #3647). Further, the raw MS data were analysed using MaxQuant (v.2.6.5.0). Protein search and identification was done using the integrated Andromeda search engine against the *Methanosarcina mazei* (DSM #3647) proteome. Only trypsin/P-specificity was considered and up to two missed cleavages, a minimal length of 7 residues, fixed Carbamidomethylation and variable methionine oxidation and N-terminal protein acetylation. Orbitrap default settings were applied. A maximum false detection rate (FDR) of 1% was used. Calculation of Label free quantification (LFQ) and intensity based total quantification (iBAQ) were enabled. Remaining parameters were kept as default settings. To semi-quantitatively determine MtrI abundance, iBAQ values for MtrI and MtrA where divided and plotted as bar-plots. P-values were calculated using a paired t-test in GraphPad Prism (v.10) of log_2_-transformed iBAQ values. For His_6_-MtrI pulldowns, iBAQ values of triplicates were ranked according to their mean-values and plotted against their rank. Plots were made in GraphPad Prism (v.10).

### Molecular size determination

We performed size exclusion chromatography (SEC) by injecting 400 μl of DDM-solubilised, purified protein into a Superose 6 Increase 10/300 GL column (Cytiva), previously equilibrated with buffer A and 0.05% DDM and attached to an Äkta pure system. Chromatography was performed at 4°C and a flow rate of 0.4 ml/min. Protein-elution was monitored by UV-absorbance at 280 nm. To analyse the molecular weight of the protein complex by mass photometry a OneMP mass photometer was used and data was acquired with AcquireMP v.2.3 (Refeyn). As mass photometry of Mtr was impossible in the presence of detergent-micelles, we used Mtr complex that was solubilised with LMNG – but washed and eluted without detergent. Movies were recorded at 1 kHz, with exposure times ranging from 0.6 to 0.9 ms, to optimize signal while avoiding saturation. Glass slides were cleaned with isopropanol and Mili-Q water and silicon gaskets were placed onto the clean slides. Following instrument calibration, 19 μl of buffer A was pipetted into a gasket well before finding the focus. The focal position was locked using the autofocus function of the instrument. The measurement started after adding and mixing 1 μl of 0.5 μM protein onto the gasket well. Data analysis was performed using DiscoverMP software (Refeyn).

### Cryo-EM and grid preparation

Grids for cryo-EM of the Mtr complex were prepared using QUANTIFOIL R 2/1 copper 300 mesh grids (Quantifoil Micro Tools). Prior to sample application, the grids were glow-discharged for 25 seconds at 15 mA using a PELCO easiGlow device (Ted Pella). LMNG-solubilised Mtr at a concentration of 6 mg/mL was pre-mixed in a 9:1 ratio with 10 mM CHAPSO, resulting in a final CHAPSO concentration of 1 mM, to counter preferred particle orientation. Immediately after mixing, 4 μL of the protein solution was applied to the grid and plunge-frozen in liquid ethane using a Vitrobot Mark IV (Thermo Fisher Scientific). Grids were blotted for 6 seconds with a blot force of 6, while the chamber-atmosphere being kept at 4 °C and 100% humidity.

### Data collection and processing

Cryo-EM data for LMNG-solubilised Mtr was acquired using Smart EPU Software on a TFS Krios G4 cryo-TEM operating at an accelerating voltage of 300 keV and equipped with a Falcon 4i Direct Electron Detector (Thermo Fisher Scientific) and Selectrics Energy Filter set at a 10 eV slit width. Data were acquired at a nominal magnification of 165 000 x corresponding to a calibrated pixel size of 0.73 Å. Images were acquired with an exposure dose of 60 e^–^ Å^-2^ in counting mode and exported as an electron-event representation (EER) file format. A dataset containing 21 552 micrographs was acquired. The entire processing was done in CryoSPARC (Punjani *et al*., 2017). The EER files were fractionated into 80 frames, motion corrected using patch motion correction (Zheng *et al*., 2017), followed by contrast transfer function (CTF) estimation. Importing beam-shifts and successive optical grouping (38 groups) improved later estimation of higher order aberrations. Manually picking of 500-600 particles was followed by training of a Topaz model (Bepler *et al*., 2019). Topaz extract of all micrographs led to an initial particle set of 1.1 million particles. Particles were initially extracted at a box size of 450 pixels and subjected to heterogeneous refinement to sort particles in 3D. An initial map, that was generated from a testing dataset, was used three times as an input seed. One of the three heterogenous refinement output maps, containing 450 704 particles, was further re-extracted at a box size of 576 pixels and refined to 2.44 Å. Masked 3D classification without alignment (20 classes, filtered at 6 Å) was performed by placing a soft mask on the MtrA subunit. Masks were created with ChimeraX map eraser tool or the molmap command, followed by map filtering and successive CryoSPARC import. Six out of 20 classes (139 221 particles) showed MtrA bound to the membrane plane. The particles were subjected to non-uniform refinement followed by reference-based motion correction and another non-uniform refinement yielding the Mtr consensus-map at a resolution of 2.06 Å with C_1_-symmetry. To better resolve MtrA, we applied local refinement using the same MtrA-mask as in the 3D-classification, yielding a local map at a resolution of 2.49 Å. To resolve MtrH, we created a soft mask around the flexible MtrH dimer region and performed 3D variability analysis with five modes to solve at a filtered resolution of 5 Å. Successive 3D Variability Display of a single component in cluster mode led to 20 clusters filtered at 5 Å, seven of which showed MtrH in similar positions. These seven clusters, containing 225 093 particles were further subjected to masked 3D classification using the same MtrH mask. Out of four classes, one (56 229 particles) was locally refined to obtain a map of 3.2 Å resolution. To create a complete Mtr composite map, MtrH was further C_3_-symmetrized using Volume alignment tools with 120° 3D rotations. The composite map was constructed in ChimeraX using the vop max command upon normalizing the maps to the same threshold with vop scale. The final composite incorporated the consensus map, the locally refined MtrA and the C_3_-symmetrized MtrH.

### Model building and refinement

Initial models were built with AlphaFold2 (Jumper *et al*., 2021). To interpret the unknown density between MtrCDE and MtrA, we used ModelAngelo (v.1.0) (Jamali *et al*., 2024) without providing a sequence and using the consensus map as an input. ModelAngelo modelled a continuous peptide of 52 residues into the density. A protein BLAST showed highest similarity to a hypothetical protein that is conserved in *Methanosarcina* species. As the protein was not annotated in the protein database, we performed *tBLASTn* to retrieve the corresponding *M. mazei* sequence. The best-matching result by far was a reading frame of 204 nucleotides encoding a 68-residue protein with the locus name MM_2401 and the UniProt accession Q8PUD4. ChimeraX and Coot (v.0.9.8.91) (Emsley *et al*., 2010) were used for manual model building. Real-space refinements of models were performed iteratively with the PHENIX-package (v.1.21-5207). Ramachandran, reference model and secondary structure restraints were applied during refinement. CIF files for ligands like the etherlipids were created using AceDRG in the CCP4 suite or eLBOW in PHENIX. Maps were graphically depicted using ChimeraX and Coot.

## Data availability

Several cryo-EM maps have been deposited in the Electron Microscopy Data Bank (EMDB) under following accession codes. Composite map: EMD-53361, Consensus map: EMD-53359, MtrA local refinement: EMD-53360, MtrH local refinement: EMD-53358. Corresponding coordinate files have been deposited in the RCSB Protein Data Bank (PDB) under following accession codes: 9QTS, 9QTQ, 9QTR, 9QTP.

The mass spectrometry proteomics data have been deposited to the ProteomeXchange Consortium (http://proteomecentral.proteomexchange.org) via the PRIDE partner repository (Vizcaíno *et al*., 2013) with the dataset identifier <PXD000XXX>."

## Acknowledgements

We acknowledge the contributions of the cryo-EM Facility of the Philipps-University Marburg. We thank Rolf Thauer for continuous support and helpful discussions. We thank Darja Deobald from the UFZ Leipzig for performing metal quantification through ICP–MS. We thank Georg Hochberg from the University of Marburg for enabling access to mass photometry. This work was supported by the Deutsche Forschungsgemeinschaft Emmy Noether grant (SCHU 3364/1-1, to JMS), RTG 2937 (to JMS) and SCHM1052/20-2 (to RAS), as well as the European Union’s Horizon 2020 research and innovation programme (Two-CO_2_-One; grant agreement no. 101075992 to JMS. The views and opinions expressed are those of the author(s) only and do not necessarily reflect those of the European Union or the European Research Council. Neither the European Union nor the granting authority can be held responsible for them. T.R-T. acknowledges the funding from the International Max Planck Research School Principles of Microbial Life.

## Author contributions

**Tristan Reif-Trauttmansdorff**: Conceptualization, Data curation; Software; Validation; Investigation; Methodology; Writing—original draft; Writing—review and editing. **Eva Herdering**: Data curation; Software; Validation; Investigation; Methodology; Writing—review and editing. **Stefan Bohn:** Investigation; Software; Methodology; Resources; Data curation**. Tomas Pascoa:** Investigation; Software; Data curation; Writing—review and editing. **Jörg Kahnt:** Methodology; Resources; Data curation **Erik Zimmer:** Conceptualization; Investigation; Methodology. **Anuj Kumar:** Investigation; Software; Data curation**. Ruth A. Schmitz**: Conceptualization; Resources; Data curation; Funding acquisition; Validation; Investigation; Methodology; Project administration; Writing—review and editing. **Jan M. Schuller**: Conceptualization; Resources; Data curation; Funding acquisition; Validation; Investigation; Methodology; Project administration; Writing— original draft; Writing— review and editing.

## Disclosure and competing interests statement

The authors declare no competing interests

## Notes

### Competing Interest Statement

The authors have declared no competing interest.

