## Supplemental information for "Structure of the energy converting methyltransferase (Mtr) of *Methanosarcina maze*i in complex with an oxygen-stress responsive small protein"

### Supplementary

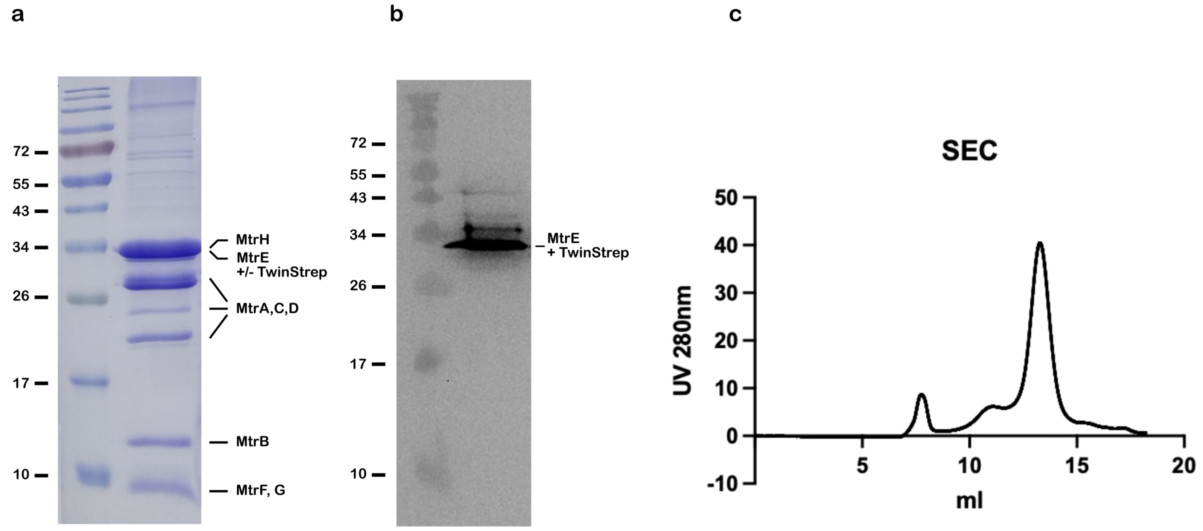

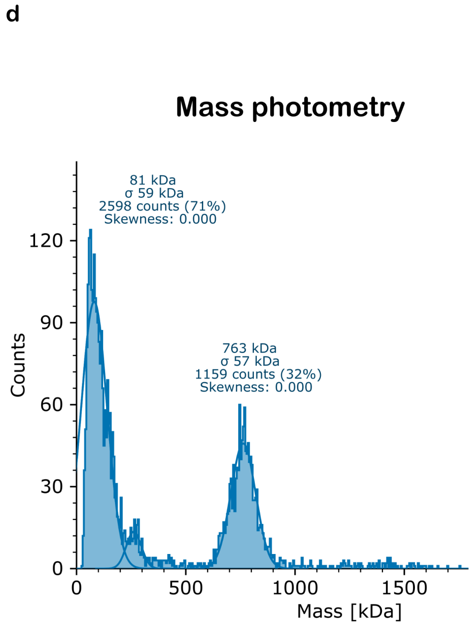

e

| # | Accession | Description | Score | Coverage | # Peptides | # PSM |
|---|---|---|---|---|---|---|
| 1 | P80650 | MtrH | 3028,63 | 80,38% | 31 | 1740 |
| 2 | Q03001 | Dystonin OS=Homo sapiens | 0 | 65,68% | 568 | 1360 |
| 3 | O59640 | **MtrA** | 1117,75 | 89,17% | 24 | 868 |
| 4 | P11532 | Dystrophin OS=Homo sapiens | 0 | 71,45% | 310 | 746 |
| 5 | P15924 | Desmoplakin OS=Homo sapiens | 17,95 | 77,99% | 327 | 741 |
| 6 | P80651 | MtrE | 694,16 | 21,71% | 10 | 670 |
| 7 | Q8PW00 | Rubrerythrin OS=Methanosarcina mazei | 702,46 | 95,40% | 36 | 591 |
| 8 | P04264 | Keratin, type II cytoskeletal 1 OS=Homo sapiens | 507,69 | 75,62% | 75 | 560 |
| 9 | P80655 | MtrB | 1076,19 | 57,41% | 7 | 553 |
| 10 | Q07283 | Trichohyalin OS=Homo sapiens | 0 | 80,44% | 255 | 551 |
| 11 | P80656 | MtrG | 1087,48 | 98,61% | 12 | 545 |
| 12 | Q60186 | V-type ATP synthase alpha chain OS=Methanosarcina mazei | 306,62 | 75,43% | 46 | 516 |
| 47 | Q8PW06 | 60 kDa chaperonin OS=Methanosarcina mazei | 149,03 | 87,31% | 52 | 195 |
| 48 | P80654 | MtrF | 355,66 | 50,00% | 5 | 191 |
| 49 | Q8PTW8 | RAP domain-containing protein OS=Methanosarcina mazei | 0 | 50,84% | 101 | 187 |
| 594 | Q8PXC3 | Probable RNA 2'-phosphotransferase OS=Methanosarcina mazei | 5,87 | 63,29% | 19 | 40 |
| 595 | O59638 | MtrC | 44,47 | 12,36% | 4 | 40 |
| 596 | Q8PSK9 | PPK2 domain-containing protein OS=Methanosarcina mazei | 0 | 57,35% | 20 | 40 |
| 996 | Q8PT68 | ABC transporter, permease protein OS=Methanosarcina mazei | 0 | 20,25% | 10 | 16 |
| 997 | P80653 | MtrD | 9,07 | 4,00% | 2 | 16 |

**Fig. S1. Mtr-purification and protein analysis. a**, 15% SDS-gel of concentrated StrepTactin eluate. **b**, SDS-gel corresponding anti-TwinStrep-tag antibody western blot. **c**, Size-exclusion chromatography of DDM-solubilised Mtr complex using a Superose 6 10/300 increase column. Elution at 13.3 ml corresponds to a complex size as previously reported for Mtr. **d**, mass photometry of LMNG-solubilised and successively detergent-freed Mtr complex. **e**, mass spectrometry results of LMNG-solubilised and purified Mtr complex, analysed with ProteomeDiscoverer.

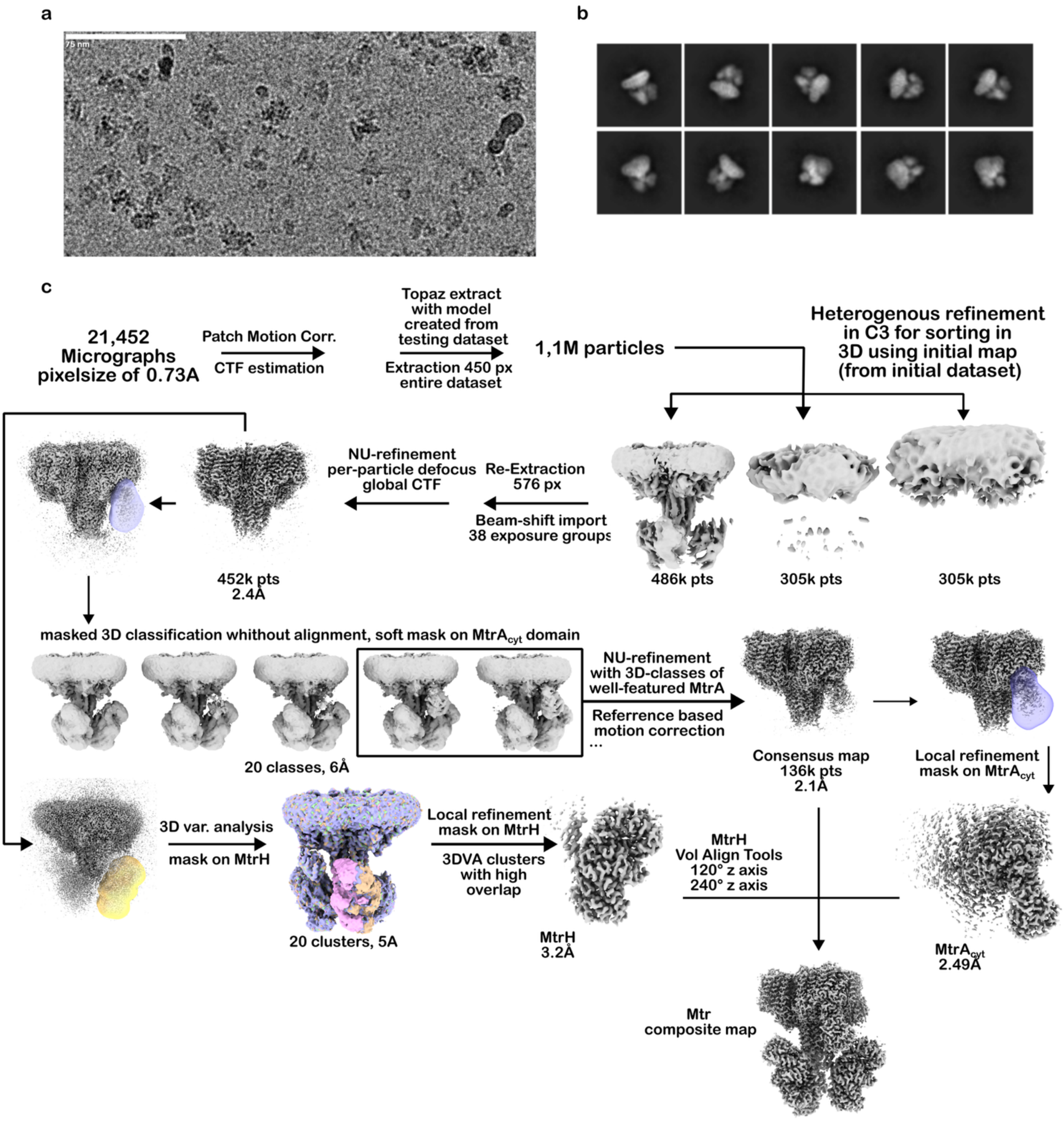

**Fig. S2. CryoEM data processing workflow. a,** Exemplary micrograph with 75 nm scale bar. **b**, Selected 2D class averages depicting diverse particle orientations. **c**, Mtr data processing tree.

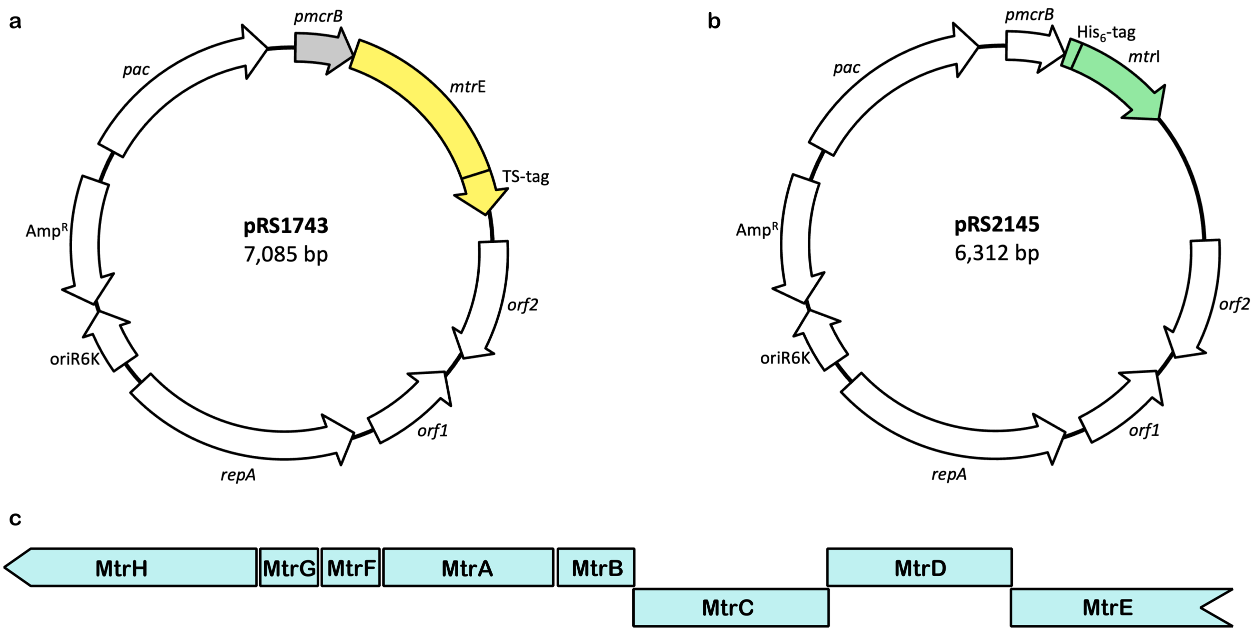

d

MKCEACGRESDTKYCNDCGKVMDEVVRRVGEARWAAIDDCSFIYPLVQRVGRGEATVNDIIQALDVED

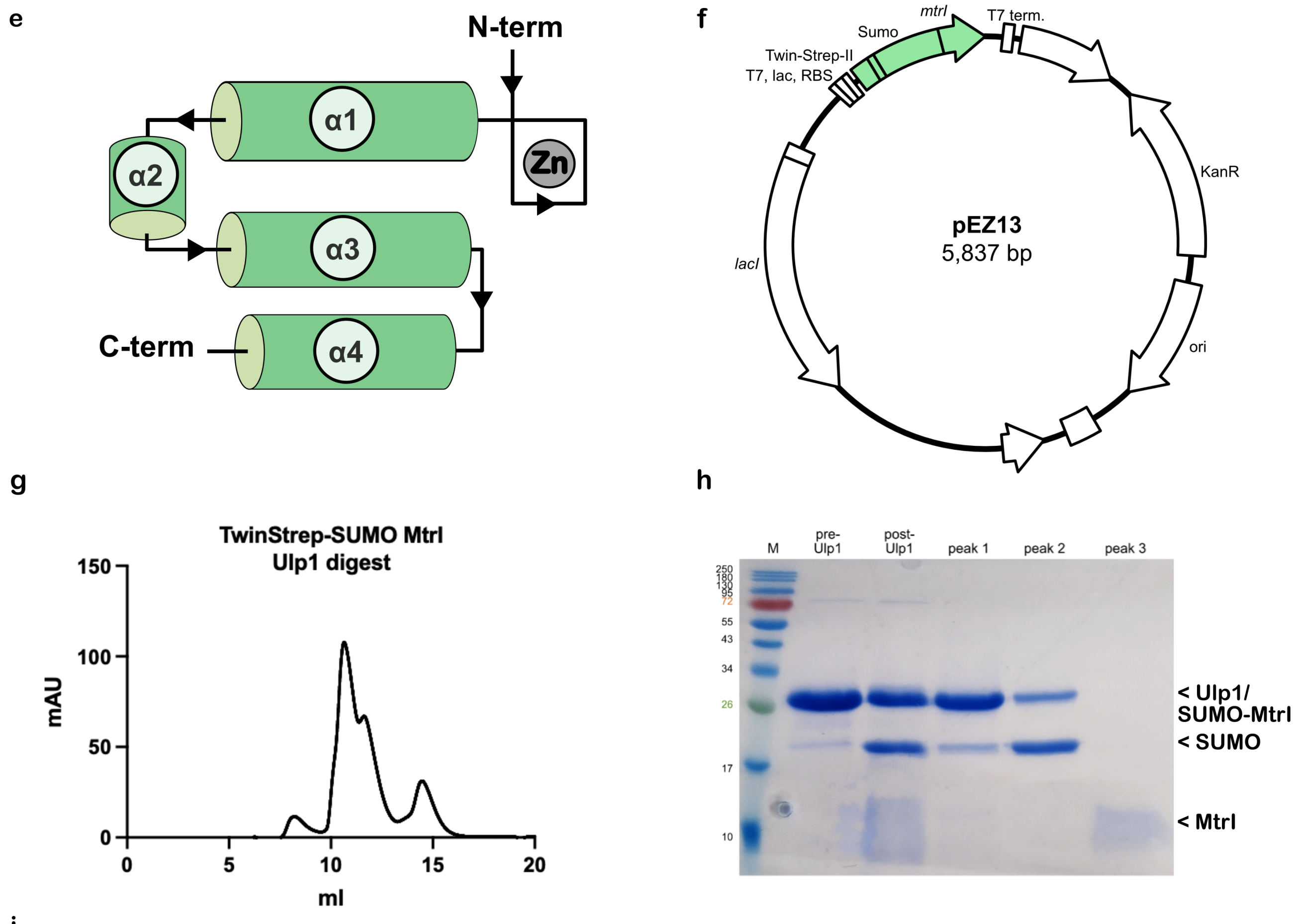

i

| Sample | $^{56}$Fe / Protein | ±$^{56}$Fe/Protein | $^{59}$Co / Protein | ±$^{59}$Co/Protein | $^{66}$Zn / Protein | ±$^{66}$Zn/Protein |
|---|---|---|---|---|---|---|
| MtrI_SEC_peak3 | <0 | 0,02 | 0,00 | 0,00 | 0,90 | 0,04 |

**Fig. S3. Plasmids and additional information on MtrI. a**, Map of plasmid pRS1743 used for expression of MtrE with a C-terminal TS-tag in *M. mazei*. Shown in white are the pRS1595 backbone elements (Thomsen and Schmitz 2022), in grey the *pmcr*B promotor and in yellow the *mtr*E-gene with TS-tag. **b**, Map of plasmid pRS2145 used for expression of MtrI with a N-terminal $His_6$-tag in *M. mazei*. Shown in white are the pRS1807 backbone elements (Hüttermann and Schmitz 2024) and in green the *mtrI*-gene and $His_6$-tag. **c**, Operon structure of Mtr in *M. mazei*. **d,** protein sequence of MM_2401 (MtrI). **e,** Secondary structure topology of MtrI. **f,** Map of plasmid pEZ13 used for recombinant expression of TS-SUMO-MtrI in *E. coli* BL21. **g**, Size-exclusion-chromatography of concentrated StrepTactin-eluates of TS-SUMO-MtrI on a Superdex 75 Increase 10/300 column. Double-peak at around 10-12 ml correspond to TS-SUMO-MtrI and TS-SUMO. Peak at around 14.5 ml correspond to MtrI cleavage product. **h**, SDS-gel of the TS-SUMO-MtrI purification. Lanes from left to right: Protein ladder, StrepTactin-eluate before Ulp1-digest, StrepTactin-eluate after Ulp1-digest, SEC-peak at 10.7 ml, SEC-peak at 11.7 ml, SEC peak at 14.5 ml. **I**, ICP-MS of purified MtrI measured as technical triplicates. Values are calculated based on a predicted molecular weight of 7.62 kDa.

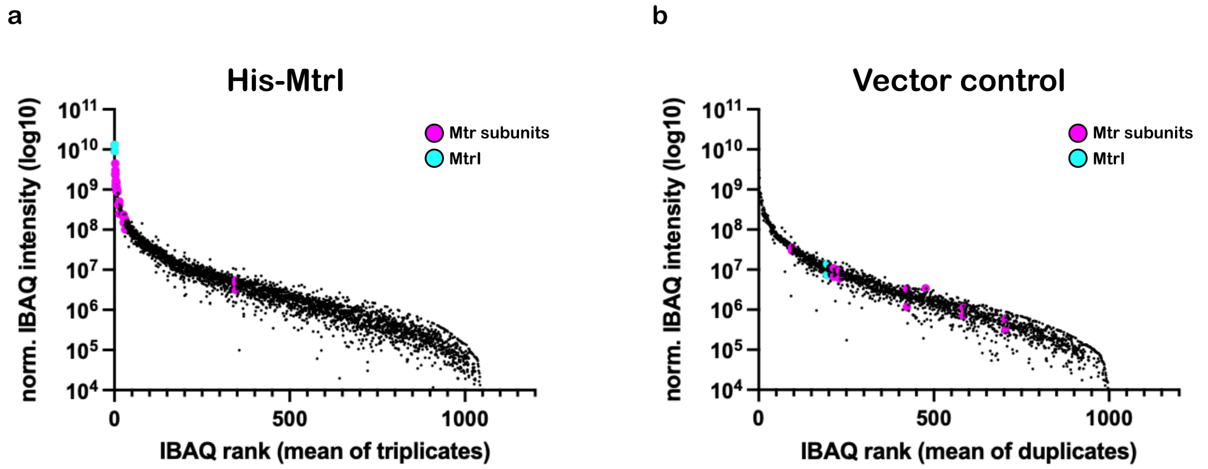

**Fig. S4. Label-free quantification of Mtr subunits by His$_6$-MtrI pulldown. a**, Mass spectrometry analysis of Ni-NTA purification from *M. mazei* expressing His$_6$-tagged MtrI. $Log_{10}$(iBAQ) values of proteins (triplicates, calculated with MaxQuant v.2.6.5.0) are plotted against their rank position (mean of triplicates). **b**, Equivalent analysis from an empty-vector *M. mazei* control strain shown as $log_{10}$(iBAQ) values (duplicates) plotted against their rank position. **a, b**, His$_6$-MtrI pulldown leads to clear enrichment of Mtr subunits compared to the empty-vector control. The reciprocal pulldown of Mtr subunits confirms the stable association of MtrI with the complex.

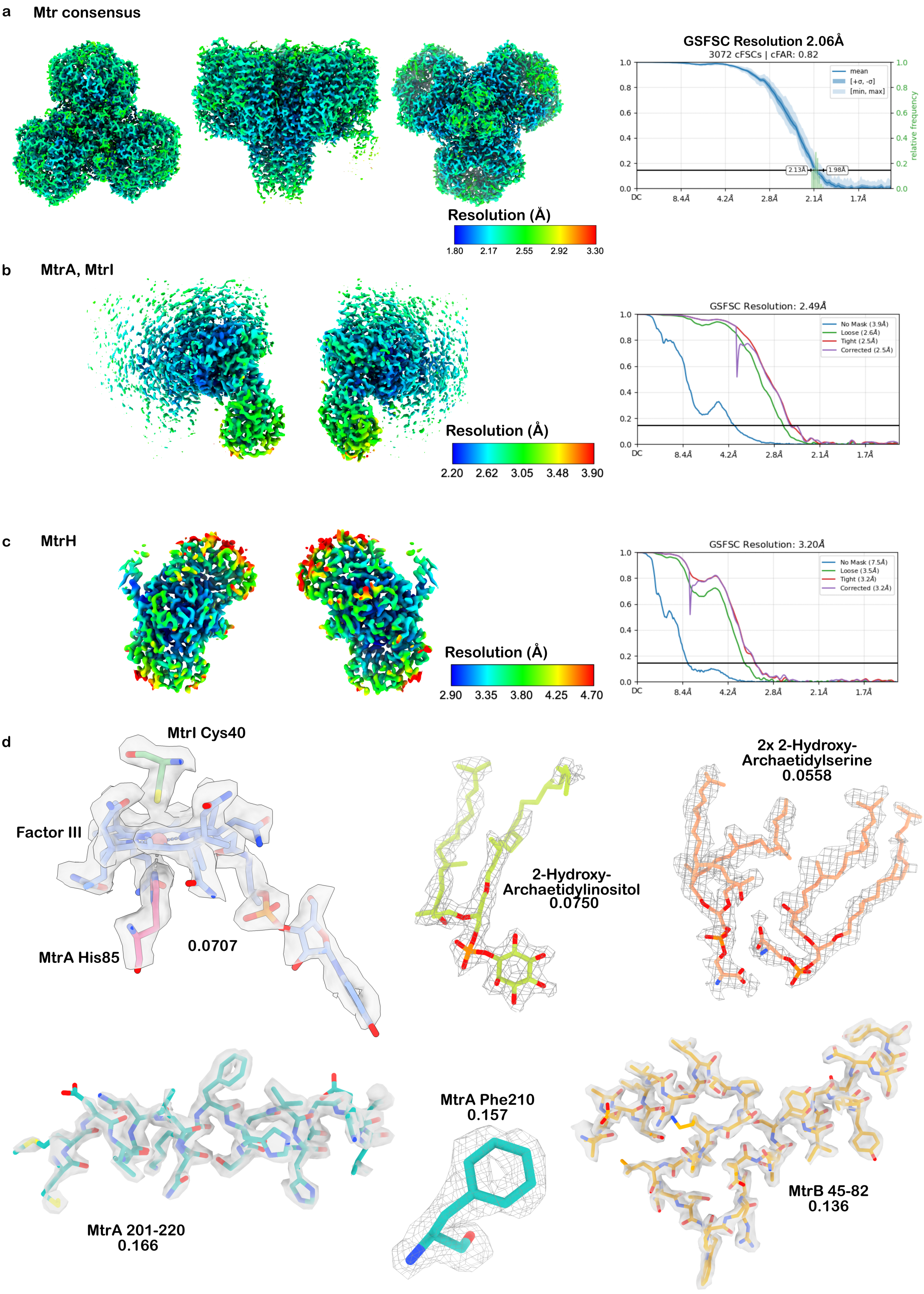
a
Mtr consensus
GSFSC Resolution 2.06Å
3072 cFSCs | cFAR: 0.82
mean
[+σ, -σ]
[min, max]
relative frequency
2.13Å
1.98Å
DC
8.4Å
4.2Å
2.8Å
2.1Å
1.7Å
Resolution (Å)
1.80 2.17 2.55 2.92 3.30
b
MtrA, MtrI
GSFSC Resolution: 2.49Å
No Mask (3.9Å)
Loose (2.6Å)
Tight (2.5Å)
Corrected (2.5Å)
Resolution (Å)
2.20 2.62 3.05 3.48 3.90
c
MtrH
GSFSC Resolution: 3.20Å
No Mask (7.5Å)
Loose (3.5Å)
Tight (3.2Å)
Corrected (3.2Å)
Resolution (Å)
2.90 3.35 3.80 4.25 4.70
d
MtrI Cys40
Factor III
MtrA His85
0.0707
2-Hydroxy-Archaetidylinositol
0.0750
2x 2-Hydroxy-Archaetidylserine
0.0558
MtrA 201-220
0.166
MtrA Phe210
0.157
MtrB 45-82
0.136

**Fig. S5. Local resolution estimates and quality of cryo-EM-maps. a**-**c**, Cryo-EM maps of Mtr consensus (core), locally refined MtrA and locally refined MtrH colored by local resolution values estimated in cryoSPARC**.** Gold standard Fourier shell correlation (GSFSC) plots show resolutions at a 0.143 threshold (black line). **d**, Representative regions of the Cryo-EM maps shown at corresponding threshold levels together with atomic models displayed as sticks, demonstrate the high quality of the maps.

### Supplementary data table 1, CryoEM data collection, refinement and validation

| | **Mtr Composite map** (EMDB-53361) (PDB: 9QTS) | **MtrA local map** (EMDB-53360) (PDB:9QTR) | **Mtr consensus map** (EMDB-53359) (PDB:9QTQ) | **MtrH local map** (EMDB-53358) (PDB:9QTP) |
|---|---|---|---|---|
| **Data collection and processing** | | | | |
| Magnification | 165 000x | 165 000x | 165 000x | 165 000x |
| Voltage (kV) | 300 | 300 | 300 | 300 |
| Electron exposure (e–/Å$^2$) | 60 | 60 | 60 | 60 |
| Defocus range (µm) | 0.5-2 | 0.5-2 | 0.5-2 | 0.5-2 |
| Pixel size (Å) | 0.73 | 0.73 | 0.73 | 0.73 |
| Symmetry imposed | C1 | C1 | C1 | C1 |
| Initial particle images (no.) | 1 096 091 | 1 096 091 | 1 096 091 | 1 096 091 |
| Final particle images (no.) | 174 856 | 136 531 | 136 531 | 56 229 |
| Map resolution (Å) | | 2.49 | 2.49 | 2.33 |
| FSC threshold | | 0.143 | 0.143 | 0.143 |
| Map resolution range (Å) | | 2.2-3.2 | 1.8-2.8 | 2.9-3.8 |
| **Refinement** | | | | |
| Initial model used (PDB code) | *de novo,* ModelAngelo, AlphaFold | *de novo,* ModelAngelo, *AlphaFold* | *de novo,* ModelAngelo, *AlphaFold* | De novo, AlphaFold |
| Model resolution (Å) | | 2.8 | 2.1 | 3.3 |
| FSC threshold | | 0.5 | 0.5 | 0.5 |
| Model resolution range (Å) | | 2.4-2.8 | 2.0-2.1 | 3.1-3.3 |
| Map sharpening *B* factor (Å$^2$) | | 58 | 38.3 | 62.6 |
| Model composition | | | | |
| Non-hydrogen | 45429 | 1698 | 28656 | 5082 |
| atoms | 5432 | 215 | 3231 | 670 |
| Protein residues | 19 | 1 | 18 | 0 |
| Ligands | | | | |
| *B* factors (Å$^2$) | | | | |
| Protein | 42.58 | 31.70 | 29.12 | 75.51 |
| Ligand | 53.32 | 14.60 | 35.59 | |
| R.m.s. deviations | | | | |
| Bond lengths (Å) | 0.002 | 0.003 | 0.005 | 0.003 |
| Bond angles (°) | 0.456 | 0.676 | 0.634 | 0.563 |
| Validation | | | | |
| MolProbity score | 2.13 | 2.65 | 1.83 | 2.06 |
| Clashscore | 22.44 | 9.16 | 13.81 | 7.59 |
| Poor rotamers (%) | 1.46 | 6.98 | 1.66 | 2.57 |
| Ramachandran plot | | | | |
| Favored (%) | 97.08 | 91.00 | 97.93 | 95.17 |
| Allowed (%) | 2.87 | 8.53 | 2.04 | 4.68 |
| Disalowed (%) | 0.04 | 0.47 | 0.03 | 0.15 |
